# GeST: Towards Building A Generative Pretrained Transformer for Learning Cellular Spatial Context

**DOI:** 10.1101/2025.04.09.648072

**Authors:** Minsheng Hao, Nan Yan, Haiyang Bian, Yixin Chen, Jin Gu, Lei Wei, Xuegong Zhang

## Abstract

Learning spatial context of cells through pretraining on spatial transcriptomics (ST) data may empower us to systematically decipher tissue organization and cellular interactions. Yet, transformer-based generative models often focus on modeling individual cells, neglecting the intricate spatial relationships within them. We develop GeST, a deep transformer model that is pretrained by a novel spatially informed generation task: Predict cellular expression profile of a given location based on the information from its neighboring cells. We propose a spatial attention mechanism for efficient pretraining, a flexible serialization strategy for converting ST data into sequences, and a cell tokenization method for quantizing gene expression profiles. We pretrained GeST on large-scale ST datasets of different ST technologies and demonstrated its superior performance in generating unseen spatial cells. Our results also show that GeST can extract spatial niche embeddings in a zero-shot way and can be further fine-tuned for spatial annotation tasks. Furthermore, GeST can simulate gene expression changes in response to perturbations of cells within spatial context, closely matching existing experimental results. Overall, GeST offers a powerful generative pre-training framework for learning spatial contexts in spatial transcriptomics.

## 1 Introduction

In recent years, transformer-based models pre-trained on large-scale scientific data have emerged as a new paradigm in artificial intelligence for biology [Webb and others, 2018; Bunne *et al*., 2024; Szałata *et al*., 2024], allowing the development of foundation models tailored to specific modalities such as DNA sequences [Nguyen *et al*., 2024], proteins [Abramson *et al*., 2024] and single cell gene expression [Theodoris et al., 2023; Hao et al., 2024; Cui *et al*., 2024; Bian *et al*., 2024]. However, most of these models focus on gene-gene relationships within isolated cellular contexts, neglecting the intricate cell-cell communications in spatial scenarios. As a result, current models struggle to handle spatial tasks such as understanding spatial cell patterns, which limits their ability to fully comprehend cellular behaviors in complex tissue systems.

Spatial transcriptomics (ST) is a rapidly developing technology that combines high-throughput gene expression profiling with spatial localization of cells within tissue sections [Moses and Pachter, 2022]. Utilizing ST to understand the spatial context of cells is critical for deciphering tissue organization mechanisms [Palla *et al*., 2022; Wu *et al*., 2021] and has great value in biological and medical research like identification of therapeutic biomarkers [Zhang *et al*., 2022]. Previous studies such as SpaGCN [Hu *et al*., 2021] and GraphST [Long *et al*., 2023] often applied graph neural networks to integrate gene expression and spatial information to learn spatial representations. These models were trained independently for each dataset, leaving the paradigms of pretraining or generative modeling unexplored. Currently, rapidly growing ST datasets enable us to learn cell-cell relationships in a data-driven manner. Beyond single-cell transcriptomics, where a cell is analogous to a sentence composed of gene tokens, in spatial transcriptomics, a tissue is a document consisting of many cell sentences. A recent study called CellPLM [Wen *et al*., 2023] built a BERT-style [Devlin, 2018] pre-trained model by using partial gene expression data from a target cell and information from its neighboring cells to predict the remaining gene expression. However, since CellPLM needs to know the expression of a subset of genes before predicting the cell’s complete gene expres-sion, it cannot generate brand new cells in unseen locations. This limitation restricts its ability to explicitly study how spatial context alone influences the characteristics of a cell. In addition, constrained by the BERT modeling, its predictions are based on the existing input all at once, lacking the ability to adapt to dynamic spatial contexts, further hindering various downstream applications such as *in-silico* spatial perturbation.

Inspired by the advancement of GPT models [Achiam *et al*., 2023; Radford *et al*., 2019; Brown, 2020], we endeavor to develop a generative pre-trained transformer tailored to ST data to overcome the above limitations and support new applications. We propose a spatially informed generation task for ST data, in which the model’s objective is to iteratively generate cells’ expression at given positions based on information of their neighboring cells (Figure 1). However, such modeling of ST data faces several unique challenges. First, unlike regular grid image data, ST data are known for their irregular data structure, where cells are distributed arbitrarily in 2-dimensional tissue sections with varying numbers. Methods like Vision Transformer [Dosovitskiy, 2020], which consider 2D data as a sequence with a fixed number and order, are not suitable for the generation task of ST data. Therefore, in order to unlock the full potential of irregular spatial structure, it is critical to design a special attention mechanism for our spatially informed generation task together with a flexible serialization strategy. Second, directly generating continuous gene expression values of cells may introduce error accumulation during the autoregressive inference [Pasini *et al*., 2024]. Thus, we need a robust tokenization method to map cells into discrete tokens, like what we do in natural language processing.

**Figure 1:**
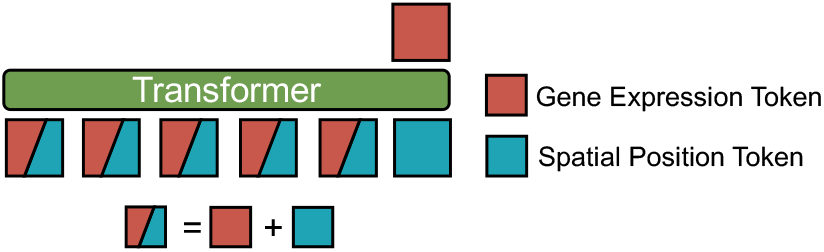
Spatially informed generation task: Given the spatial location *x* of one target cell, the transformer model takes its spatial coordinate *s*(*x*), its neighbors’ *N* (*x*) gene expression *g*(*N* (*x*)) and spatial location *s*(*N* (*x*)) as the input, and predicts the target cell’s gene expression *g*(*x*).

To address these challenges, we present **GeST**, a deep **Ge**nerative pre-trained transformer for **ST** data which generates cells by leveraging the neighbor information. After pre-training on a large corpus of ST data, our model can learn biologically meaningful spatial patterns and can also be fine-tuned for other applications. Moreover, it can explore perturbation effects in spatial contexts by manipulating the given neighborhood information. To the best of our knowledge, GeST is the first generative pre-trained transformer to understand cell-cell relationships and advance cell modeling in the spatial context. Taken together, our work makes the following key contributions:

- **Spatially Informed Generation of ST Data**: We develop novel modeling for generative pre-trained transformers on ST data. To implement this task with high computational efficiency and flexibility, we design an attention mechanism called *Spatial Attention* and a serialization strategy to convert ST data into a sequence.
- **Cell Tokenization Method**: We develop a cell tokenization method to quantize cells’ expression profiles to discrete tokens, along with a hierarchical pre-training loss designed to mitigate error accumulation in autoregressive generation.
- **Superior performance in Downstream Tasks**: We conducted several experiments and comparisons to show GeST superior performance in generating unseen spatial cells, spatial clustering and annotation tasks.
- **Spatial Perturbation Analysis**: We establish GeST as a pioneering model for *in-silico* spatial perturbation analysis. The simulation results align with real experiments, which provide an *in-silico* extension of current single-cell perturbation studies.

## 2 Related Work

### 2.1 Transformer models for transcriptomics data

The recent broad applications of transformer-based models have inspired researchers to develop specialized methods for transcriptomics data [Szałata *et al*., 2024]. In 2022, scBERT [Yang *et al*., 2022] uses a BERT-style transformer architecture to extract embeddings of single-cell transcriptomics data by masked token prediction, and fine-tunes the model for cell type annotation. Theodoris *et al*.[2023] propose Geneformer, to learn gene-gene network dynamics from self-attention mechanisms by pretaining it on a large-scale corpus of single-cell transcriptomcis data, and further use it to *in-silico* perturbation analysis. scGPT [Cui *et al*., 2024], scFoundation [Hao *et al*., 2024] and scMulan [Bian *et al*., 2024] are foundation models built for single-cell transcriptomics with millions of parameters in transformer models. After pretraining, they are applied to diverse downstream tasks, such as cell clustering, supervised cell type annotation, and batch correction, all achieving superior performance. However, these transformer-based methods only focus on isolated cell modeling due to the lack of spatial information in single-cell transcriptomics data. Recently, ST datasets have increased rapidly, and a few studies have begun to leverage them for pretraining. CellPLM [Wen *et al*., 2023] proposes a cell language model by considering one cell’s partial gene expression together with other cells in the input sequence to reconstruct the complete expression profile of the target cell. Still, generative modeling for ST data is under-explored. Nicheformer [Schaar *et al*., 2024] uses both ST and single-cell transcriptomics datasets for pre-training. However, it does not incorporate spatial information during the pretraining task and only designs spatially relevant fine-tuning tasks to apply to ST data.

### 2.2 Representation learning for spatial transcriptomics data

The majority of methods for learning informative representations of ST data are based on graph neural networks. SpaGCN [Hu *et al*., 2021] integrates gene expression, spatial location and histology image by a graph convolutional network to identify spatial domains in a self-supervised manner. STAGATE [Dong and Zhang, 2022] uses a graph attention auto-encoder to learn low-dimensional latent embeddings that incorporate both spatial information and gene expression profiles. GraphST [Long *et al*., 2023] adopts a contrastive learning strategy to extract representation by graph neural networks. NicheCompass [Birk *et al*., 2024] is a bio-inspired graph learning method that incorporates existing knowledge of cell-cell communication and transcriptional regulation pathways to learn an interpretable latent space of cells across multiple ST samples. However, these methods are often trained on small ST datasets due to the limited capabilities of graph neural networks. This may overlook the potential of existing large volumes of ST data.

## 3 Task Formulation

Given a spatial transcriptomics dataset, we denote it as a set {*x*_1_, *x*_2_, *x*_3_, …, *x*_*n*_}, encompassing all *n* cells within a 2-dimensional tissue slice. We define two essential functions: *g*(*·*), the gene expression retrieval function, and *s*(*·*), the spatial information retrieval function. We also define that for any given cell *x*, the set *N* (*x*) = *{x*_*N*1_, *x*_*N*2_, …, *x*_*Nk*_*}* denotes all *k* of its neighboring cells.

### Spatially informed generation (Pretraining task)

In order to learn cell-cell relationships within the spatial context, we propose a generation pretraining task aware of spatial dependency. In particular, given a target cell *x*, the objective is to predict the gene expression *g*(*x*) based on its spatial location *s*(*x*) and the information of its neighboring cells *N* (*x*), i.e., conditional distribution:

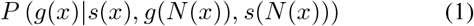

Following this modeling, the objective function of our task is

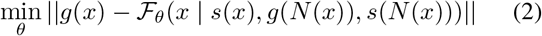

where ℱ_*θ*_ represents our proposed spatial generative model parameterized by *θ*. However, spatial data lack a natural sequential order, which challenges the application of auto-regressive models that usually work for sequence prediction tasks. To address this, we first devise a serialization strategy to convert a set of cells *N* (*x*_*k*+1_) into an ordered sequence (*x*_1_, *x*_2_, …, *x*_*k*_). Then, we can transform the objective function into a sequential format:

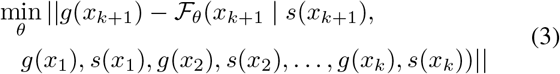

More importantly, similar to natural language generation, the objective of this task can be extended to generate multiple cells. This can be achieved by iteratively applying the function ℱ of Equation 3 to sequentially generate the gene expression of cells adjacent to the known cell neighbors.

### Niche clustering/annotation (Downstream tasks)

After the pretraining stage, we can transfer the model to diverse downstream tasks. In the studies of spatial transcriptomics, the concept ‘niche’ refers to a functional or structural tissue region where cells interact with each other and their surroundings. Identifying and understanding these niches is crucial for elucidating tissue organization [Jain and Eadon, 2024]. The objective of niche clustering or annotation task is to extract the spatial and gene expression information of a cell *x* and its neighbors *N* (*x*) into a latent embedding space that can be used for clustering or label classification. This encoding process can be formalized as follows:

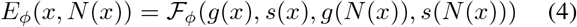

Here, *E*_*ϕ*_ represents the encoding function parameterized by *ϕ*, which integrates both the gene expression and spatial information of a cell as well as its neighbors into a unified embedding vector.

## 4 Methodology

We introduce GeST, a deep generative pretrained transformer model for learning cellular spatial context. The model has three components: an ST data serialization strategy, a cell tokenization method, and a spatial context-aware decoder. These modules together with the pretraining and fine-tuning design are detailed in the following subsections (Figure 2).

**Figure 2:**
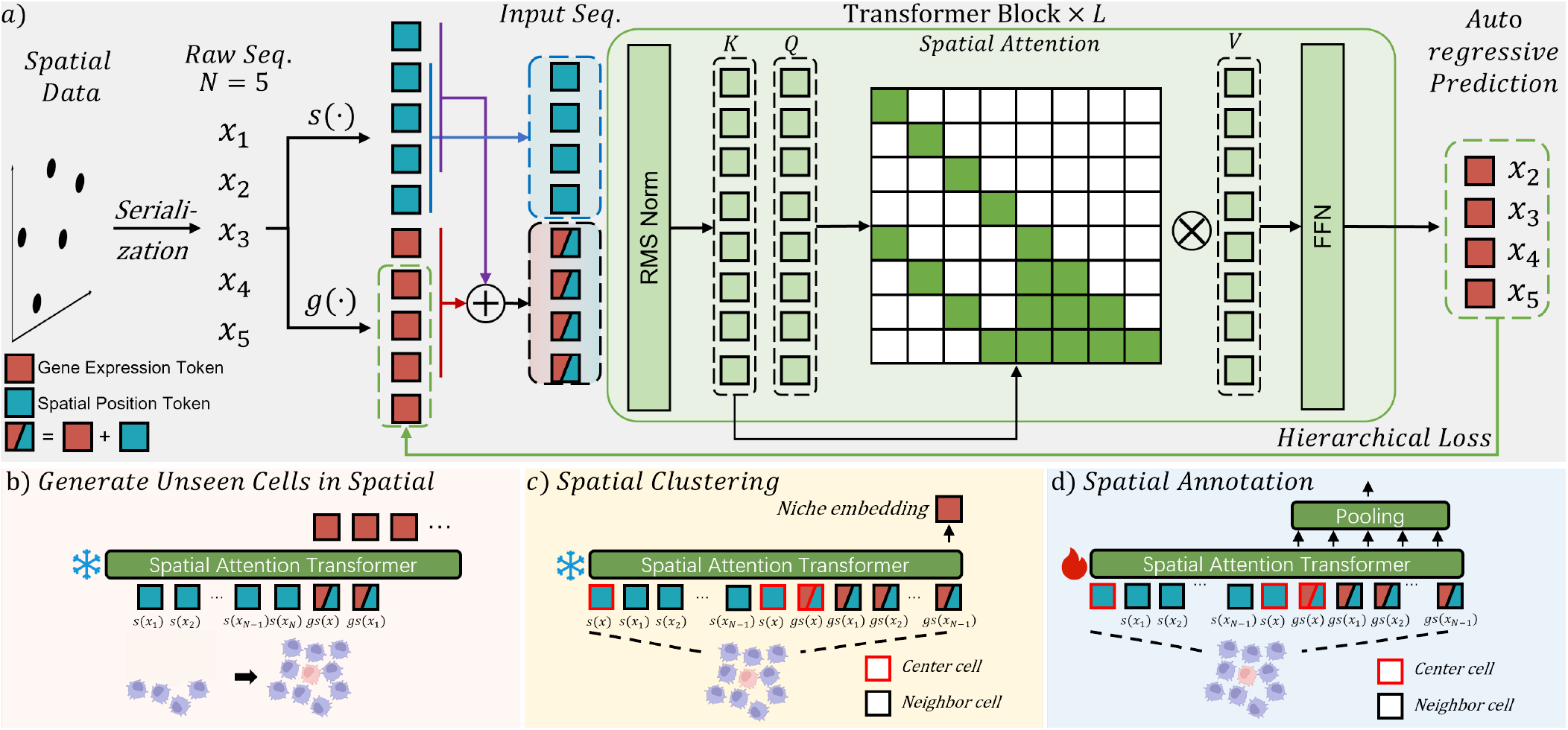
Schematic overview of GeST. a) Model architecture. b) Generating unseen cells in spatial from neighborhood cells c) Extracting a niche embedding from a group of neighborhood cells and doing spatial clustering d) Annotating spatial niches by fine-tuning GeST.

### 4.1 Spatial Context-Aware Decoder

Our main model is a transformer decoder [Vaswani, 2017] modified for the spatially informed generation task. During pre-training, the model’s input is divided into two contiguous sequences after tokenization: the neighbor cell sequence and the target cell position sequence, constituting a sequence of *N* + (*N* − 1) tokens. The output is a sequence of *N* − 1 gene expression tokens (Figure 2a).

#### Neighbor Cell Sequence

This part consists of the complete tokens of the first to the (*N* − 1)-th cells, totaling *N* − 1 cells. Each token combines both gene expression and spatial information: [*gs*(*x*_1_), *gs*(*x*_2_), …, *gs*(*x*_*N* − 1_)], where *gs*(*x*_*i*_) = *g*(*x*_*i*_) + *s*(*x*_*i*_) represents the combined embedding of gene expression and spatial position for cell *x*_*i*_.

#### Target Cell Position Sequence

This part has the spatial position tokens of the second to the *N*-th cells, totaling *N* − 1 cells, formulating a target cell position token sequence, [*s*(*x*_2_), *s*(*x*_3_), …, *s*(*x*_*N*_)].

As illustrated in Figure 2a, for each target cell position *s*(*x*_*i*+1_), we use the transformer decoder’s parallel training capability to predict the gene expression of the next neighbor cell *g*(*x*_*i*+1_). Prediction of target cell *x*_*i*+1_ is conditioned on the complete tokens of the neighbor cells {*gs*(*x*_1_), *gs*(*x*_2_), …, *gs*(*x*_*i*_)} and the spatial position token *s*(*x*_*i*+1_).

To achieve this, we design a special attention matrix called ***Spatial Attention***. Unlike the causal attention used in language models, which employs a lower triangular mask to ensure that each token can only attend to previous tokens in the sequence, our Spatial Attention allows each position to attend to specific relevant tokens, enabling the model to capture spatial dependencies more effectively. Specifically, for a sequence length of 2*L* (where *L* = *N* − 1), the attention mask *M* is a 2*L* × 2*L* matrix. For the token at position *i* + *L* (corresponding to predicting the gene expression of cell *x*_*i*+1_), we allow attention to 1) The neighbor cell tokens at positions 1 to *i* (i.e., {*gs*(*x*_1_), *gs*(*x*_2_), …, *gs*(*x*_*i*_)}). and 2) The target cell position token at position *i* + *L* (i.e., *s*(*x*_*i*+1_)). Formally, for *i* ∈ [1, *L*], the attention mask *M* is defined as:

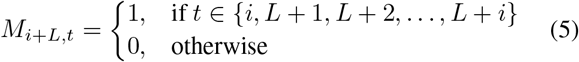

This design leverages the transformer decoder’s capability for parallel computation while effectively modeling spatial relationships [Radford *et al*., 2019]. By allowing each prediction to attend to the relevant neighbor cells and the spatial position of the target cell, the model learns to generate gene expressions conditioned on spatial context. After the decoder, a multilayer perceptron is used to convert the hidden embedding ***h*** ∈ ℝ^*D*^ to gene expression space ***ŷ*** ∈ ℝ^*T*^. Each element of prediction ***ŷ*** represents the expression level of a specific gene, where *T* is the total number of genes.

### 4.2 Serialization strategy

To get the input sequence at each training step, we crop a square in the training tissue section and use the set of all *N* cells 𝒳 = {*x*_*o*1_, *x*_*o*2_, …, *x*_*oN*_} in this square to constitute a training sequence. We serialize them by sampling along diagonal paths. Specifically, we first randomly select a cell from four vertexes of the square as the start point *x*_1_. Then we calculate all cells’ Euclidean distances from *x*_*oi*_ and use them as sampling weights: {*w*_*o*1_, *w*_*o*2_, *w*_*o*3_, …, *w*_*oN*_}, where *w*_*oi*_ = ||*p*(*x*_*oi*_) − *p*(*x*_1_) || _2_ and *p*(*x*) = (*p*_1_, *p*_2_) denotes the 2D coordinates of the cell. We do sampling without replacement for *N* − 1 times to get a sequence (*x*_1_, *x*_2_, *x*_3_, …*x*_*N*_). At each sampling time *t*, the probability of selecting cell *x*_*oi*_ is:

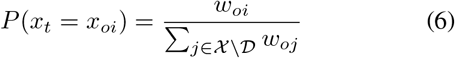

where 𝒟 contains cells that have been selected into the sequence. This strategy allows the spatially adjacent cells to have similar indexes in sequence and still retain randomness to prevent over-fitting of the order.

### 4.3 Cell tokenization

Given a cell sequence, we define the two retrieval functions *g*(*x*), *s*(*x*) to tokenize cell’s gene expression and spatial position into *d* dimension. For the spatial position *s*(*x*), we set up a coordinate system whose origin is the centered cell of the tissue section. Then we calculate the relative coordinates to the origin of other cells and perform normalization on them. We tokenize the normalized coordinate values by a 2D sinusoidal positional encoding (SPE) function SPE: ℝ^2^ → ℝ^*d*^ similar to ViT [Dosovitskiy, 2020].

For the gene expression *g*(*x*), a naive idea is to directly set the continuous cellular expression vector as the regression target. However, our preliminary experiments demonstrated that error accumulation could lead to model failure during the autoregressive generation process (refer to “w/o quantization” in Table 4). Therefore, we develop a “meta cell vocabulary” 𝒞 to quantize cells’ continuous expression to discrete tokens. Different from gene bin strategy proposed by previous work, we quantize the whole cell into one token which is derived from the skeleton of the data.

Given a training dataset with *n* cells and *T* genes, we perform PCA reduction to *p* dimensions and group all cells into *K* clusters by K-means. Then the center point of each cluster is a “meta cell”. Meta cells capture the skeleton of the data and are used as the basic units of the meta cell vocabulary. Each meta cell *c*_*i*_ has a mean PCA vector 𝒞_pca_[*i*] and a mean gene expression vector 𝒞_expr_[*i*] of its corresponding cluster.

For cell *x* in the sequence, we get its gene expression PCA reduction *x*_*pca*_, find the index of its nearest meta cell in the PCA space and take the meta cell’s expression as the input token to the model, which can be formulated as:

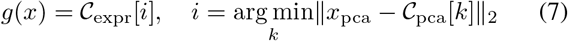

### 4.4 Loss function

To compute the loss between the predicted ***ŷ*** and the ground truth gene expression token *g*(*x*), we propose a hierarchical cross-entropy loss function. We note that our quantization strategy (see Sec4.3) could lose a continuous distance relationship in the expression space. Therefore, we further perform the K-means algorithm on the meta cell vocabulary with a small number of clusters to obtain hierarchical labels at different levels. For a unified notation, a meta cell *c* is associated with hierarchical labels at four levels: *l*_0_(*c*), *l*_1_(*c*), *l*_2_(*c*), and *l*_3_(*c*), each of them has *K, K*_1_, *K*_2_, and *K*_3_ categories, where *l*_0_(*c*), *K* represent the initial fine-grained meta cell vocabulary. Then we project the model outputs ***ŷ*** to logits **z** ∈ ℝ^*K*^ corresponding to each meta cell:

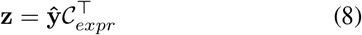

where 𝒞_*expr*_ ∈ ℝ^*K* ×*T*^ contains the meta cells’ expression profiles. We then apply the softmax function for normalization. The predicted probability that the prediction ***ŷ*** belongs to meta cell *c* is:

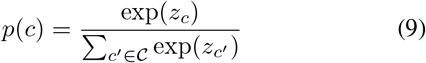

To compute the hierarchical losses, we aggregate probabilities across hierarchical levels. For level *i*, the probability of label *k* is calculated as:

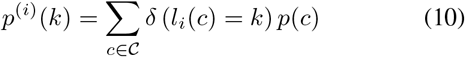

where *δ*(*·*) is the Kronecker delta function, which equals 1 if the condition is true and 0 otherwise.

The overall loss function ℒ is defined as a weighted sum of the negative log-likelihood losses at each hierarchical level:

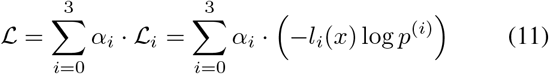

where *α*_*i*_ are weights and we set 0.25 as default. ℒ_*i*_ is the cross-entropy loss at level *i*, and *l*_*i*_(*x*) is the ground truth label of the target cell’s nearest meta cell at level *i*. By minimizing this hierarchical loss function, the model is encouraged to make correct predictions at multiple levels, making it robust to single level wrong predictions.

With regard to the inference strategy (Figure 2b), there are two modes to convert the predicted probability to the final output gene expression value vector: (1) “Picking” mode: we directly use the meta cell’s expression with the highest probability as the prediction. (2) “Weighted aggregation” mode: we set *p*(*c*) as the weight to aggregate all meta cells’ expression as the prediction.

### 4.5 Niche embedding extraction

After pretraining, GeST can be used for niche clustering in a zero-shot manner and can be fine-tuned for niche annotation. Both tasks require the extraction of a niche embedding. Given a niche comprising a target cell *x* and its *N* − 1 neighboring cells, we first take both position tokens and cell tokens as input. Then, we include the target cell’s position token *s*(*x*) twice: once at the beginning and once at the end of all position tokens, resulting in a total of *N* + 1 position tokens. For the cell sequence, we incorporate the content tokens of all cells, yielding *N* tokens (Figure 2c). Under this configuration, the last output token of the model is generated based on the information from all cells, and we define this as the niche embedding, consistent with Equation 4. We use the hidden embedding ***h*** for the zero-shot niche clustering task.

For fine-tuning, we input the sequence in the same setting and further apply a mean pooling operation to aggregate these embeddings into a single vector (Figure 2d) ***h***_*p*_ = Pool({***h***_*x*_} ∪ {***h***_*n*_ |*n* ∈ *N* (*x*)}). The embedding ***h***_*p*_ is then passed through a linear layer to obtain the logits for the niche classification task.

## 5 Experiments

We conducted four experiments for demonstration: unseen cell generation, zero-shot niche embedding clustering, fine-tuning-based niche label annotation, and *in-silico* spatial perturbation. Sec5.1 shows our model’s superior performance in the spatial generation task. Sec5.2 and Sec5.3 show the generalization ability to other downstream applications. Sec5.4 is to explore a cutting-edge analysis method of *in-silico* perturbation studies in biology.

### 5.1 Unseen cell generation

To validate the generative capability of our model, we used three datasets from different ST technologies and resolutions covering normal and disease samples: a single-cell resolution MERFISH dataset of the whole mouse brain [Zhang *et al*., 2023], a multi-cell resolution Visium dataset of human primary liver cancer (PLC) [Wu *et al*., 2021], and a sub-cell resolution Stereo-seq dataset of the mouse brain sagittal section [Cheng *et al*., 2022]. For each dataset, we divided all tissue sections into training, validation, and testing sets (see supplementary for detailed descriptions). We cropped a region of the section in the testing set and named it as an “unseen region”. Our task is to predict the gene expression of this unseen region based on the remaining part of the section. Since there are no existing methods designed for spatial generation of ST data, we trained two models as baselines: a gaussian process (GP) model and a multilayer perceptron (MLP). These two models used cells’ absolute spatial coordinates and gene expressions from the uncropped areas in each section as training data.

Recognizing that not all genes exhibit strong spatial patterns, we also focused on the top 200 spatially variable genes (SVGs) per slide, identified using SOMDE [Hao *et al*., 2021]. We calculated three evaluation metrics for each testing set (Table 1). Our model in the “picking” mode (labeled as “Ours”) achieved lower root mean square error (RMSE) of both all genes and top 200 SVGs compared to the baseline models. Switched to the “weighted aggregation” mode (labeled as “Ours-W”), our model produced even lower regression error in the MERFISH dataset and consistently surpassed baselines in the Spearman’s coefficient. In a detailed analysis of one section in the testing set of the MERFISH dataset (Figure 3), our model more accurately predicted genes with higher spatial variation, a trend less evident in the baseline models. These findings highlight our model’s capability of learning the underlying spatial characteristics of gene expression and cell organization by the spatially informed generation task.

**Table 1:** Benchmark on unseen cell generation. R.all: Root mean square error (RMSE) of all genes. R.200: Root mean square error of top 200 SVGs. *ρ*: Spearman’s rank correlation coefficient. The best scores of each testing set are in **bold**, and the second best are in <u>underline</u>.

| Method | MERFISH |  |  |  |  |  |  |  |  | Visium |  |  | Stereo-seq |  |  |
| --- | --- | --- | --- | --- | --- | --- | --- | --- | --- | --- | --- | --- | --- | --- | --- |
|  | Anterior Brain |  |  | Mid Brain |  |  | Posterior Brain |  |  | Primary Liver Cancer |  |  | Sagittal Brain |  |  |
| | R.all↓ | R.200↓ | $\rho$ ↑ | R.all↓ | R.200↓ | $\rho$ ↑ | R.all↓ | R.200↓ | $\rho$ ↑ | R.all↓ | R.200↓ | $\rho$ ↑ | R.all↓ | R.200↓ | $\rho$ ↑ |
| Ours | 1.374 | 1.296 | 0.301 | 1.379 | 1.319 | 0.265 | 1.386 | 1.340 | 0.242 | **1.303** | **1.126** | **0.540** | 1.405 | 1.357 | **0.326** |
| Ours-W | **1.352** | **1.244** | **0.340** | **1.367** | **1.288** | **0.302** | **1.376** | **1.322** | **0.275** | 1.320 | 1.150 | 0.499 | **1.399** | **1.340** | 0.323 |
| GP | 1.379 | 1.289 | 0.241 | 1.389 | 1.337 | 0.241 | 1.393 | 1.354 | 0.214 | 1.357 | 1.264 | 0.272 | 1.413 | 1.405 | 0.073 |
| MLP | 1.399 | 1.369 | 0.319 | 1.404 | 1.393 | 0.283 | 1.406 | 1.397 | 0.267 | 1.347 | 1.174 | 0.491 | 1.403 | 1.357 | 0.314 |

**Figure 3:**
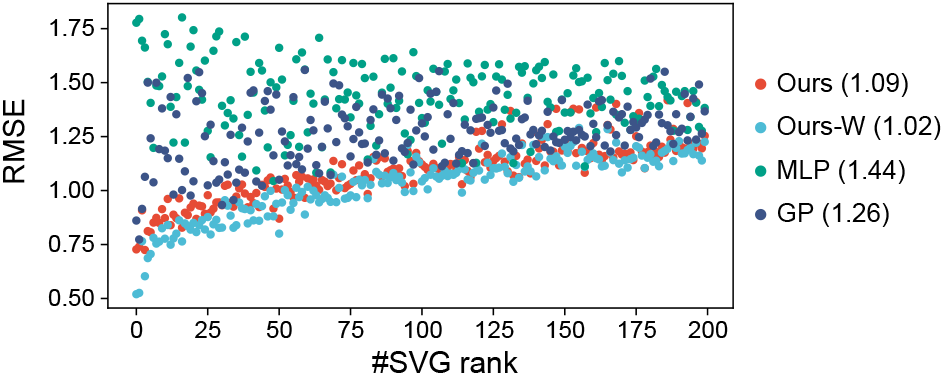
Benchmark on top 200 spatially variable genes (SVGs) expression prediction. Each dot represents a gene. The x-axis is the rank of spatial variation, higher rank representing higher spatial variation. The number in parentheses is the mean RMSE of all these SVGs.

### 5.2 Unsupervised niche clustering

As a pretrained model, one key feature of GeST is its ability to generalize to various downstream applications. We evaluated this capability on spatial niche clustering and annotation tasks in the MERFISH mouse brain dataset, as it is well-annotated with two levels of anatomical labels, “Division” and “Region”, according to the Mouse Brain Common Coordinate Framework (CCF) v3 [Wang *et al*., 2020]. For the niche clustering task, we used niche-level embeddings of each cell for clustering. We compared our method with four methods: NicheCompass, GraphST, STAGATE and SpaGCN. We also included a baseline that gets cluster labels based solely on the cell’s own gene expression without spatial information (Raw). As shown in the Table 2, all spatial clustering methods except SpaGCN outperformed the raw baseline on the adjusted mutual information (AMI) scores. Our pretrained model (labeled as “Ours”) achieved higher AMI than other graph-based methods in most test sets. As expected, after we continued training our model on the test data in the same generative way (labeled as “Ours-FT”), the model showed consistently higher performance. These results demonstrate that our pre-trained model can be effectively transferred to new tissues in a zero-shot manner.

**Table 2:** Benchmark of AMI scores on unsupervised niche clustering tasks. NicheC: NicheCompass. The best scores of each testing set are in **bold**, and the second best are in <u>underline</u>.

| Method | Anterior Brain |  | Mid Brain |  | Posterior Brain |  |
| --- | --- | --- | --- | --- | --- | --- |
|  | Division | Region | Division | Region | Division | Region |
| Ours | 0.344 | **0.406** | 0.550 | 0.535 | 0.460 | 0.472 |
| Ours-FT | **0.368** | 0.404 | **0.585** | **0.589** | **0.489** | **0.528** |
| GraphST | 0.220 | 0.361 | 0.485 | 0.463 | 0.405 | 0.365 |
| NicheC | 0.253 | 0.386 | 0.547 | 0.551 | 0.449 | 0.440 |
| STAGATE | 0.229 | 0.382 | 0.527 | 0.516 | 0.447 | 0.449 |
| SpaGCN | 0.126 | 0.191 | 0.244 | 0.265 | 0.207 | 0.196 |
| Raw | 0.083 | 0.183 | 0.233 | 0.288 | 0.216 | 0.219 |

### 5.3 Supervised Niche annotation

Currently, spatial annotation methods are lacking due to limited data with fine annotation. Therefore, we compared two single cell annotation methods: scANVI [Xu *et al*., 2021] and Celltypist [Domínguez Conde *et al*., 2022], which only utilize gene expression of one cell for classification. As detailed in Table 3, our model greatly outperformed the single-cell methods at both resolutions. This is because the niche labels were annotated in terms of both gene expression and spatial location information of every cell, and our model perceives all the cells in their spatial context rather than treating them individually.

**Table 3:** Benchmark of F1 score on supervised niche annotation task. The best scores of each testing set are in **bold**.

| Method | Anterior Brain |  | Mid Brain |  | Posterior Brain |  |
| --- | --- | --- | --- | --- | --- | --- |
|  | Division | Region | Division | Region | Division | Region |
| Ours | **0.711** | **0.483** | **0.549** | **0.392** | **0.462** | **0.312** |
| scANVI | 0.309 | 0.222 | 0.337 | 0.216 | 0.264 | 0.126 |
| CellTypist | 0.106 | 0.032 | 0.205 | 0.066 | 0.131 | 0.042 |

**Table 4:** Ablation study. ‘Ours’ represents the default model with 8 layers and 8 heads (L8H8) trained on full data and 600µm window size. ‘800µm+L’ is a model with L12H8 trained on 800µm window size. ‘w/o spatial info’ stands for replacing all positional embedding with an all-ones vector. R.all: RMSE of all genes. R.50: RMSE of top 50 SVGs. *ρ*: Spearman’s rank correlation coefficient.

|  | Ours | Model Size |  |  | Data |  | Window Size |  |  | w/o hierarchy | w/o quantization | w/o spatial info | random order |
| --- | --- | --- | --- | --- | --- | --- | --- | --- | --- | --- | --- | --- | --- |
| | | L2H2 | L4H4 | L16H16 | 1/2 | 1/3 | 200μm | 800μm | 800μm+L | | | | |
| R.all↓ | 1.367 | 1.371 | 1.373 | 1.362 | 1.381 | 1.375 | 1.376 | 1.371 | 1.362 | 1.382 | 1.389 | 1.397 | 1.384 |
| R.50↓ | 1.214 | 1.243 | 1.249 | 1.208 | 1.261 | 1.246 | 1.251 | 1.232 | 1.204 | 1.270 | 1.315 | 1.325 | 1.256 |
| $\rho \uparrow$ | 0.29 | 0.288 | 0.289 | 0.291 | 0.289 | 0.288 | 0.275 | 0.285 | 0.292 | 0.288 | N.A. | 0.230 | 0.287 |

### 5.4 In-silico spatial perturbation

Measuring the cell response by perturbations like diseases or treatments is of great significance in biological and medical research [Rood *et al*., 2024]. Inspired by single-cell pre-trained models [Theodoris *et al*., 2023], we establish GeST as a pioneer model for predicting cell response of *in-silico* perturbation in spatial transcriptomics. Here, we demonstrated an *in-silico* spatial perturbation experiment of an ischemic condition of the mouse brain (Figure 4a). Recently, Han *et al*. [2024] measured gene expression and cell distribution in the ischemic mouse brain and identified several ischemic regions, including the infarct core area (ICA) and the proximal region of the peri-infarct area (PIA P). In our experiment, we selected an area with a similar location of ICA from one sample as the region of interest (ROI) and generated the surrounding cell expression as the control group, representing the normal gene expression around the ROI. In order to simulate the ischemic effect, we manually altered gene expression in this ROI according to the differentially expressed genes (DEGs) of ICA. We then fed the model with the same surrounding positions and obtained the gene expression prediction as the perturbation group, representing the predicted PIA P.

**Figure 4:**
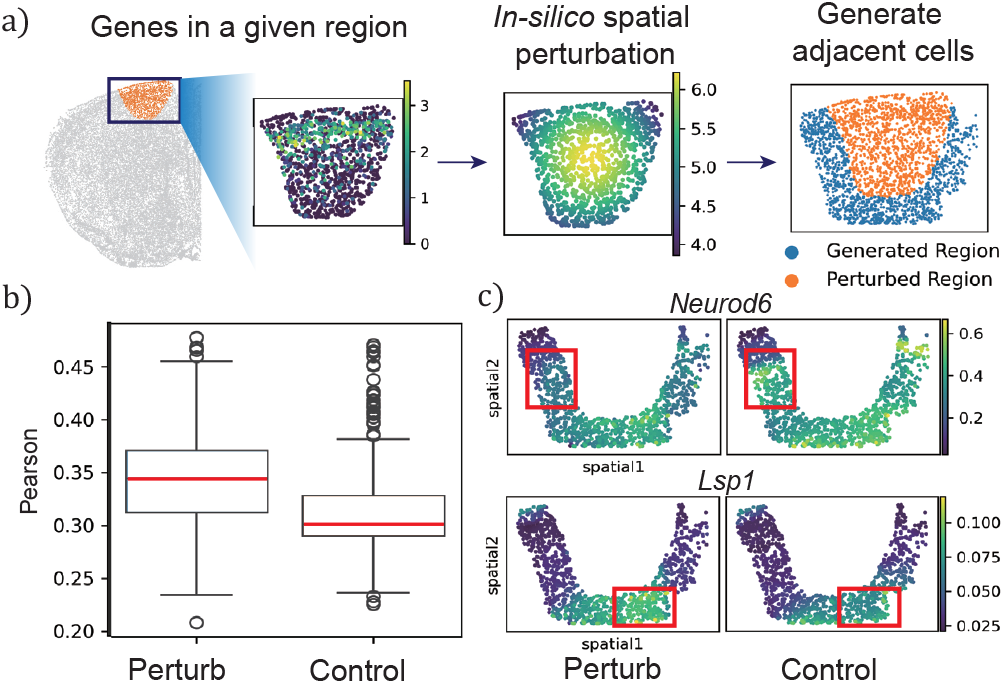
*In-silico* spatial perturbation experiment. a) Flow chart of *in-silico* spatial perturbation experiment. b) Box plot showing PCCs between perturbation group and control group. *P* -value *<* 0.001, t-test. c) Visualization of two predicted DEGs.

To validate the results, we calculated the Pearson correlation coefficients (PCCs) between gene expression of the perturbation group and the average expression of PIA P, and compared them with those between control group and PIA P. The PCCs are significantly higher in the perturbation group (Figure 4b). Taking all 87 high and low DEGs in PIA P as ground truth, we correctly classified 70.11% of them by our *in-silico* perturbation experiment. This is higher than the baseline accuracy of 44.8%, which is obtained by simply adopting the DEGs from ICA (i.e. a naive model that believes changes in ROI are the same in the neighbor). Figure 4c gives two predicted DEG examples. These findings are consistent with the experimental results from [Han *et al*., 2024].

### 5.5 Ablation study

We conducted comprehensive ablation experiments to examine the impact of model size, training data volume, window size, and the designed modules on model performance (Table 4). Increasing the model from a small one to our baseline resulted in significant performance improvements. Beyond our default model, further increases in model size yielded diminishing benefits, suggesting that our default model strikes an optimal balance between performance and computational efficiency. On the other hand, models trained on larger datasets performed better, suggesting that increasing the amount of training data could further enhance model effectiveness in future work. Neighbor window size is another important factor that controls the information density and the sequence length of the input. The model with window sizes of 600µm and 800µm achieved higher performance than that of 200µm (Table 4). We also noted that the 800µm window size didn’t introduce a higher performance, so we trained a model with a larger size (L12H8) and found that it achieved the highest performance. These results illustrated that a larger window size may introduce too much spatial variation, making the small model hard to learn. And our current model and data setting is a trade off between them.

We also ablated the hierarchy loss and cell tokenization module. We found that the model without hierarchy loss achieved a bad performance on all three metrics. For quantization module ablation, we trained a model with the mean square error (MSE) loss. We noted this MSE model generated invalid negative expression values in prediction and showed a quite low performance in RMSE, eventually causing failure in computing Spearman correlation. For spatial information, we removed all neighbor information and observed a significant performance drop. Finally, we compared our spatial ordinal serialization strategy with a random sampling strategy and observed a substantial improvement from our strategy. It was intuitively aligned with our assumption that the ordinal strategy meets the actual testing scenario and thus brings a gain in performance.

## 6 Conclusion

In this work, we present GeST, a deep generative transformer model pretrained by a novel spatially informed generation task. To the best of our knowledge, GeST is the first generative pretrained model in spatial transcriptomics. GeST features a spatial attention mechanism, paired with a serialization strategy and cell tokenization module, to model ST data in an auto-regressive generative pretraining. Our results demonstrate that our spatial generation task enables the model to learn underlying spatial contexts, thereby enhancing performance on niche-level tasks. Furthermore, we apply GeST to explore perturbation effects in spatial transcriptomics, laying the groundwork for building more comprehensive foundation models for spatial biology.

## References

[Abramson et al., 2024] Josh Abramson, Jonas Adler, Jack Dunger, Richard Evans, Tim Green, Alexander Pritzel, Olaf Ronneberger, Lindsay Willmore, Andrew J Ballard, Joshua Bambrick, et al. Accurate structure prediction of biomolecular interactions with alphafold 3. Nature, pages 1–3, 2024.

[Achiam et al., 2023] Josh Achiam, Steven Adler, Sandhini Agarwal, Lama Ahmad, Ilge Akkaya, Florencia Leoni Aleman, Diogo Almeida, Janko Altenschmidt, Sam Altman, Shyamal Anadkat, et al. Gpt-4 technical report. arXiv preprint 2303.08774, 2023.

[Bian et al., 2024] Haiyang Bian, Yixin Chen, Xiaomin Dong, Chen Li, Minsheng Hao, Sijie Chen, Jinyi Hu, Maosong Sun, Lei Wei, and Xuegong Zhang. scmulan: a multitask generative pre-trained language model for single-cell analysis. In International Conference on Research in Computational Molecular Biology, pages 479–482. Springer, 2024.

[Birk et al., 2024] Sebastian Birk, Irene Bonafonte-Pardàs, Adib Miraki Feriz, Adam Boxall, Eneritz Agirre, Fani Memi, Anna Maguza, Rong Fan, Gonçalo Castelo-Branco, Omer Ali Bayraktar, et al. Large-scale characterization of cell niches in spatial atlases using bio-inspired graph learning. bioRxiv, pages 2024–02, 2024.

[Brown, 2020] Tom B Brown. Language models are fewshot learners. arXiv preprint 2005.14165, 2020.

[Bunne et al., 2024] Charlotte Bunne, Yusuf Roohani, Yanay Rosen, Ankit Gupta, Xikun Zhang, Marcel Roed, Theo Alexandrov, Mohammed AlQuraishi, Patricia Brennan, Daniel B Burkhardt, et al. How to build the virtual cell with artificial intelligence: Priorities and opportunities. arXiv preprint 2409.11654, 2024.

[Cheng et al., 2022] Mengnan Cheng, Liang Wu, Lei Han, Xin Huang, Yiwei Lai, Jiangshan Xu, Shuai Wang, Mei Li, Huiwen Zheng, Weimin Feng, et al. A cellular resolution spatial transcriptomic landscape of the medial structures in postnatal mouse brain. Frontiers in Cell and Developmental Biology, 10:878346, 2022.

[Cui et al., 2024] Haotian Cui, Chloe Wang, Hassaan Maan, Kuan Pang, Fengning Luo, Nan Duan, and Bo Wang. scgpt: toward building a foundation model for single-cell multi-omics using generative ai. Nature Methods, pages 1–11, 2024.

[Devlin, 2018] Jacob Devlin. Bert: Pre-training of deep bidirectional transformers for language understanding. arXiv preprint 1810.04805, 2018.

[Domínguez Conde et al., 2022] C Domínguez Conde, C Xu, LB Jarvis, DB Rainbow, SB Wells, T Gomes, SK Howlett, O Suchanek, K Polanski, HW King, et al. Cross-tissue immune cell analysis reveals tissue-specific features in humans. Science, 376(6594):eabl5197, 2022.

[Dong and Zhang, 2022] Kangning Dong and Shihua Zhang. Deciphering spatial domains from spatially resolved transcriptomics with an adaptive graph attention auto-encoder. Nature communications, 13(1):1739, 2022.

[Dosovitskiy, 2020] Alexey Dosovitskiy. An image is worth 16×16 words: Transformers for image recognition at scale. arXiv preprint 2010.11929, 2020.

[Han et al., 2024] Bing Han, Shunheng Zhou, Yuan Zhang, Sina Chen, Wen Xi, Chenchen Liu, Xu Zhou, Mengqin Yuan, Xiaoyu Yu, Lu Li, et al. Integrating spatial and single-cell transcriptomics to characterize the molecular and cellular architecture of the ischemic mouse brain. Science Translational Medicine, 16(733):eadg1323, 2024.

[Hao et al., 2021] Minsheng Hao, Kui Hua, and Xuegong Zhang. Somde: a scalable method for identifying spatially variable genes with self-organizing map. Bioinformatics, 37(23):4392–4398, 2021.

[Hao et al., 2024] Minsheng Hao, Jing Gong, Xin Zeng, Chiming Liu, Yucheng Guo, Xingyi Cheng, Taifeng Wang, Jianzhu Ma, Xuegong Zhang, and Le Song. Large-scale foundation model on single-cell transcriptomics. Nature Methods, pages 1–11, 2024.

[Hu et al., 2021] Jian Hu, Xiangjie Li, Kyle Coleman, Amelia Schroeder, Nan Ma, David J Irwin, Edward B Lee, Russell T Shinohara, and Mingyao Li. Spagcn: Integrating gene expression, spatial location and histology to identify spatial domains and spatially variable genes by graph convolutional network. Nature methods, 18(11):1342–1351, 2021.

[Jain and Eadon, 2024] Sanjay Jain and Michael T Eadon. Spatial transcriptomics in health and disease. Nature Reviews Nephrology, pages 1–13, 2024.

[Long et al., 2023] Yahui Long, Kok Siong Ang, Mengwei Li, Kian Long Kelvin Chong, Raman Sethi, Chengwei Zhong, Hang Xu, Zhiwei Ong, Karishma Sachaphibulkij, Ao Chen, et al. Spatially informed clustering, integration, and deconvolution of spatial transcriptomics with graphst. Nature Communications, 14(1):1155, 2023.

[Moses and Pachter, 2022] Lambda Moses and Lior Pachter. Museum of spatial transcriptomics. Nature methods, 19(5):534–546, 2022.

[Nguyen et al., 2024] Eric Nguyen, Michael Poli, Matthew G Durrant, Armin W Thomas, Brian Kang, Jeremy Sullivan, Madelena Y Ng, Ashley Lewis, Aman Patel, Aaron Lou, et al. Sequence modeling and design from molecular to genome scale with evo. BioRxiv, pages 2024–02, 2024.

[Palla et al., 2022] Giovanni Palla, David S Fischer, Aviv Regev, and Fabian J Theis. Spatial components of molecular tissue biology. Nature Biotechnology, 40(3):308–318, 2022.

[Pasini et al., 2024] Marco Pasini, Javier Nistal, Stefan Lattner, and George Fazekas. Continuous autoregressive models with noise augmentation avoid error accumulation. arXiv preprint 2411.18447, 2024.

[Radford et al., 2019] Alec Radford, Jeffrey Wu, Rewon Child, David Luan, Dario Amodei, Ilya Sutskever, et al. Language models are unsupervised multitask learners. OpenAI blog, 1(8):9, 2019.

[Rood et al., 2024] Jennifer E Rood, Anna Hupalowska, and Aviv Regev. Toward a foundation model of causal cell and tissue biology with a perturbation cell and tissue atlas. Cell, 187(17):4520–4545, 2024.

[Schaar et al., 2024] Anna C Schaar, Alejandro Tejada-Lapuerta, Giovanni Palla, Robert Gutgesell, Lennard Halle, Mariia Minaeva, Larsen Vornholz, Leander Dony, Francesca Drummer, Mojtaba Bahrami, et al. Nicheformer: a foundation model for single-cell and spatial omics. bioRxiv, pages 2024–04, 2024.

[Szałata et al., 2024] Artur Szałata, Karin Hrovatin, Sören Becker, Alejandro Tejada-Lapuerta, Haotian Cui, Bo Wang, and Fabian J Theis. Transformers in single-cell omics: a review and new perspectives. Nature Methods, 21(8):1430–1443, 2024.

[Theodoris et al., 2023] Christina V Theodoris, Ling Xiao, Anant Chopra, Mark D Chaffin, Zeina R Al Sayed, Matthew C Hill, Helene Mantineo, Elizabeth M Brydon, Zexian Zeng, X Shirley Liu, et al. Transfer learning enables predictions in network biology. Nature, 618(7965):616–624, 2023.

[Vaswani, 2017] A Vaswani. Attention is all you need. Advances in Neural Information Processing Systems, 2017.

[Wang et al., 2020] Quanxin Wang, Song-Lin Ding, Yang Li, Josh Royall, David Feng, Phil Lesnar, Nile Graddis, Maitham Naeemi, Benjamin Facer, Anh Ho, et al. The allen mouse brain common coordinate framework: a 3d reference atlas. Cell, 181(4):936–953, 2020.

[Webb and others, 2018] Sarah Webb et al. Deep learning for biology. Nature, 554(7693):555–557, 2018.

[Wen et al., 2023] Hongzhi Wen, Wenzhuo Tang, Xinnan Dai, Jiayuan Ding, Wei Jin, Yuying Xie, and Jiliang Tang. Cellplm: pre-training of cell language model beyond single cells. bioRxiv, pages 2023–10, 2023.

[Wu et al., 2021] Rui Wu, Wenbo Guo, Xinyao Qiu, Shicheng Wang, Chengjun Sui, Qiuyu Lian, Jianmin Wu, Yiran Shan, Zhao Yang, Shuai Yang, et al. Comprehensive analysis of spatial architecture in primary liver cancer. Science Advances, 7(51):eabg3750, 2021.

[Xu et al., 2021] Chenling Xu, Romain Lopez, Edouard Mehlman, Jeffrey Regier, Michael I Jordan, and Nir Yosef. Probabilistic harmonization and annotation of single-cell transcriptomics data with deep generative models. Molecular systems biology, 17(1):e9620, 2021.

[Yang et al., 2022] Fan Yang, Wenchuan Wang, Fang Wang, Yuan Fang, Duyu Tang, Junzhou Huang, Hui Lu, and Jianhua Yao. scbert as a large-scale pretrained deep language model for cell type annotation of single-cell rna-seq data. Nature Machine Intelligence, 4(10):852–866, 2022.

[Zhang et al., 2022] Linlin Zhang, Dongsheng Chen, Dongli Song, Xiaoxia Liu, Yanan Zhang, Xun Xu, and Xiangdong Wang. Clinical and translational values of spatial transcriptomics. Signal Transduction and Targeted Therapy, 7(1):111, 2022.

[Zhang et al., 2023] Meng Zhang, Xingjie Pan, Won Jung, Aaron R Halpern, Stephen W Eichhorn, Zhiyun Lei, Limor Cohen, Kimberly A Smith, Bosiljka Tasic, Zizhen Yao, et al. Molecularly defined and spatially resolved cell atlas of the whole mouse brain. Nature, 624(7991):343– 354, 2023.

