## Supplemental Figures for "GeST: Towards Building A Generative Pretrained Transformer for Learning Cellular Spatial Context"

### 1 DATASET AND TASK DESCRIPTION

The original MERFISH spatial transcriptomics dataset has two coronal and two sagittal sliced adult mouse brain replicates with 1,122 genes, containing 2.8, 1.2, 1.6 and 0.16 million cells, respectively. All these data are mapped to the whole mouse brain taxonomy and Allen CCFv3 and each cell has a 3-dimensional coordinate. The x and y coordinates are experimentally measured and aligned to CCF, and the z coordinates are estimated. Then the multi-tissue annotation can be obtained based on the locations. Since the spatial coordinates value is an important guidance to our model, we only used the measured x and y coordinates in our training. And we selected the first coronal replicate which has the most number of cells as the training data, and used another one as the test data. The first Mouse1 dataset encompasses 2.8 million cells across 147 coronal sections, each annotated with a panel of 1,122 genes. The Mouse2 dataset is from a separate mouse brain replicate, comprising 1.2 million cells across 66 coronal sections. These 66 coronal sections are grouped into three sets according to the location of sections in the brain: Anterior Brain, Mid Brain and Posterior Brain.

The human primary liver cancer (PLC) dataset includes five cases of hepatocellular carcinoma (HCC-1 to HCC-5), one case of intrahepatic cholangiocarcinoma (ICC-1) and one case of combined hepatocellular and cholangiocarcinoma (cHC-1), containing 84,823 spots in total. We selected one slice (HCC-1L, where L represents the leading-edge section) as the test set, and took the other 20 slices as the training set. Since the data volume of PLC by Visium is much less than the mouse brain datasets by MERFISH, we trained a GeST model with fewer layers and heads (4 transformer layers and 4 heads per layer). The slice for evaluation, HCC-1L, measured the spatial gene expression from tumor to normal tissue of one patient. We cropped an area of 100 spots containing the edge of the tumor as unseen spots (labeled as “Test”), and took all the other spots as seen spots (labeled as “Ref”) (Figure 6.3). After pretraining on 20 slices, we applied GeST to generate gene expression at the location of unseen spots based on the information of the rest seen spots. We visualized the meta-cell on the UMAP and results showed that it can well preserve the original data space (Figure 6.9).

The Stereo-seq dataset has one sagittal section from the mouse brain, with in total of 60,000 data spots. Each spot is bin50 (25 μm), a typical resolution used for analyzing stereo-seq data. We segmented the cortex region from the right top corner as the test data, and used the rest of the tissue as the training data. We train GeST with the default model size.

The mouse brain ischemic dataset is collected from the ischemic hemisphere of mice subjected to photothrombosis (experiment group) and the ipsilateral hemisphere of sham mice (control group), and is sequenced by 10X Visium spatial transcriptomics platform. Each group has four coronal sections, containing 19,777 spatial transcriptomic spots in total. The spots are annotated with anatomical brain region labels, including the normal and ischemic regions. There are 425 DEGs in the ICA and 1,263 in the PIA_P. Since GeST was pre-trained on 1,122 genes of the MERFISH dataset, we used the intersection of both dataset in the *in-silico* spatial perturbation experiment.

### 2 ITERATIVE CELL GENERATION

Our model generated unseen cells based on the information from seen cells within a given window size. If the unseen region is larger than the given window size, an iterative generation is needed (as shown in Figure 6.10). Given the spatial location of all unseen cells $X_{unseen} = \{s(x_{u1}), s(x_{u2}), ..., s(x_{uN})\}$, in each iterative round, we first find a subset of the seen cells, $X_s$ from all seen cells $X_{seen}$. Each cell in $X_s$ has at least one unseen cell in its available window. These cells will be used as the reference in this round. For each cell $x_s$ in $X_s$, we generate expression for unseen cells in its window. Relatively speaking, a subset of cells $X_{pre} = \{x_{p1}, x_{p2}, ..., x_{pM}\}$ from $X_{unseen}$ will have at least one gene expression estimation given from reference cells. For instance, we assume $x_{pi}$ has n estimations $\{g_{x1}(x_{pi}), g_{x2}(x_{pi}), ..., g_{xn}(x_{pi})\}$, where $g_{x.}$ represents a gene vector function based on cell $x.$'s window. In practice, $g_{x\cdot}(\cdot)$ is a vector where each dimension is the probability of a meta cell in the meta cell vocabulary. To get the final prediction for cell $x_{ui}$, We use the mean value of all these estimations as the final value and multiply it with the meta cell vocabulary:

$$g(x_{ui}) = \frac{1}{N}\sum_{j=1}^{N} g_{xj}(x_{ui}) \cdot \mathcal{C} \tag{1}$$

Then we update $X_{seen} = X_{seen} \cup X_{pre}$ and $X_{unseen} = X_{unseen} \setminus X_{pre}$ at the end of this round. We repeat the above process in each round until $X_{unseen}$ is empty.

### 3 2D SINUSOID POSITIONAL ENCODING

We first unified the spatial coordinates to millimeters based on the CCF information. Since the absolute values of spatial coordinates in different slices are not comparable, for each training sequence, we converted the coordinates of each cell into relative coordinates. Specifically, the coordinate value of the central cell is subtracted from each cell. Then we anchored each coordinate into integers in a fixed range. In our experiments, we use [0,200) as default. Then we use 2D sinusoidal positional encodings to encode the two-dimensional coordinates into high dimension embeddings. Our approach is inspired by the method proposed in CellPLM, which employs sinusoidal functions to encode spatial coordinates in two dimensions. The encoding for a cell located at coordinates $(x, y)$ is formulated as:

$$\begin{aligned} \mathrm{PE}_{(x,y),2i} &= \sin\left(\frac{x}{10000^{2i/d}}\right), \quad \mathrm{PE}_{(x,y),2i+1} &= \cos\left(\frac{x}{10000^{2i/d}}\right) \\ \mathrm{PE}_{(x,y),2j+d/2} &= \sin\left(\frac{y}{10000^{2j/d}}\right), \quad \mathrm{PE}_{(x,y),2j+1+d/2} &= \cos\left(\frac{y}{10000^{2j/d}}\right) \end{aligned} \tag{2}$$

where $d$ is the total dimension of the positional encoding, and $i, j \in [0, d/4)$ specify the feature dimensions. This formulation extends the original sinusoidal positional encoding used in transformers to two dimensions, capturing both horizontal and vertical spatial variations.

### 4 ALGORITHMS FOR CELL EXPRESSION QUANTIZATION

**Algorithm 1:** Construction of meta cell vocabulary.

**Data:** Spatial dataset $\mathcal{X} = \{x_1, x_2, x_3, \ldots, x_n\}$; Number of meta cells $K$; Number of hierarchical labels at different levels $K_1, K_2, K_3$

**Result:** Meta cell vocabulary $\mathcal{C}$; Hierarchical labels of each meta cell $L_1, L_2, L_3$

$\mathcal{P} \in \mathbb{R}^{n\times p} \leftarrow$ Normalize gene expression $g(\mathcal{X})$ and perform PCA reduction to $p$ dimensions.
$\mathcal{P}_{\text{label}} \in \mathbb{R}^n \leftarrow$ Calculate labels of each cell in $\mathcal{P}$ by K-means algorithm with $k$ categories.
$\mathcal{C}_{\text{pca}} \in \mathbb{R}^{K\times p} \leftarrow$ Average $\mathcal{P}$ for each cluster label in PCA space.
$\mathcal{C}_{\text{expr}} \in \mathbb{R}^{K\times T} \leftarrow$ Average $\mathcal{P}$ for each cluster label in expression space.
$L_1, L_2, L_3 \in \mathbb{R}^K \leftarrow$ Calculate labels of $\mathcal{C}_{\text{pca}}$ by K-means with $K_1, K_2, K_3$ clustering numbers.

**Algorithm 2:** Query for meta cell vocabulary.

**Input:** Query spatial expression vector $x_{\text{expr}} \in \mathbb{R}^T$
**Output:** Meta cell $\boldsymbol{c} \in \mathbb{R}^T$; Hierarchical labels of the meta cell $l_1, l_2, l_3$

$x_{\text{pca}} \leftarrow$ Project $x_{\text{expr}}$ to PCA reduction space
$i \leftarrow$ Retrieve the nearest neighbor index of $x_{\text{pca}}$ in $\mathcal{C}_{\text{pca}}$
$\boldsymbol{c} \leftarrow \mathcal{C}_{\text{expr}}[i]$
$l_1, l_2, l_3 \leftarrow L_1[i], L_2[i], L_3[i]$

### 5 Details for in-silico spatial perturbation

***In-silico* activation**: For genes that were highly expressed in ICA (e.g. *Spp1, Anxa2, Rbp1*), we used a gaussian kernel function

$$f_{\text{act}}(x, y) = a \exp\left(-\frac{(x - x_0)^2 + (y - y_0)^2}{2\sigma^2}\right)$$

to replace the original expression in the ROI, where $a$ was the maximum value of all genes, $(x_0, y_0)$ was the center coordinate of the perturbation and $\sigma$ was a hyper-parameter for controlling the rate of decay.

***In-silico* inhibition**: For genes that were lowly expressed in ICA (e.g. *Lamp5, Slc17a7, Tafa1*), we used a the similar gaussian kernel function

$$f_{\text{inh}}(x, y) = g(x, y)\left[1 - \exp\left(-\frac{(x - x_0)^2 + (y - y_0)^2}{2\sigma^2}\right)\right]$$

where $g(x, y)$ represented the original expression at position $(x, y)$, $(x_0, y_0)$ was the center coordinate of the perturbation.

### 6 Extended figures & Tables

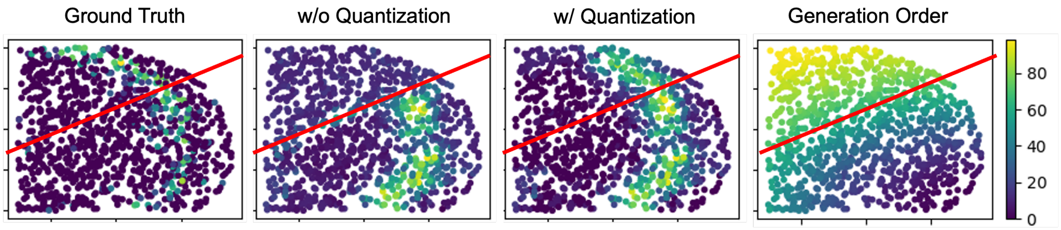


Figure 6.1: Visualizing the effect of cell quantization on multiple steps generation. Regions below the red line are reference spots, and we generated all spots above the red line by following the order shown in the right sub-figure. The color in the first three figures represents the expression value of one marker gene.

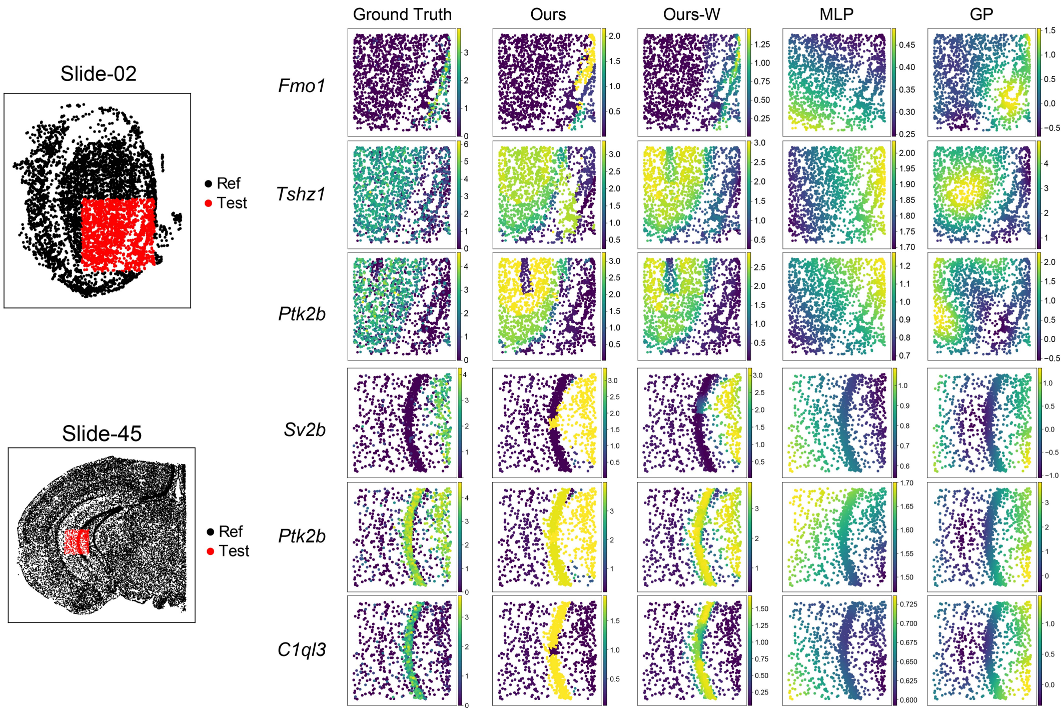


Figure 6.2: Visualization of gene expression predictions. The black and red regions indicate the reference and unseen test regions, respectively. Each row on the right shows a gene's ground truth and predicted spatial patterns.

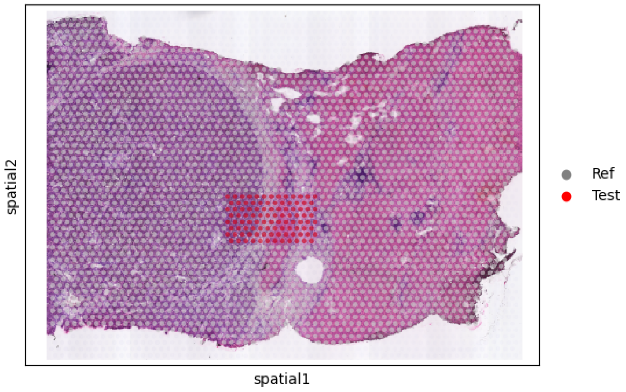


Figure 6.3: Evaluation setting of sample HCC-1L from 10X Visium PLC dataset. Left half in deep purple is tumor tissue, and the rest right half is adjacent normal tissue. 100 spots containing the edge of tumor as are labeled as 'Test' set, and all the other spots are labeled as 'Ref' set.

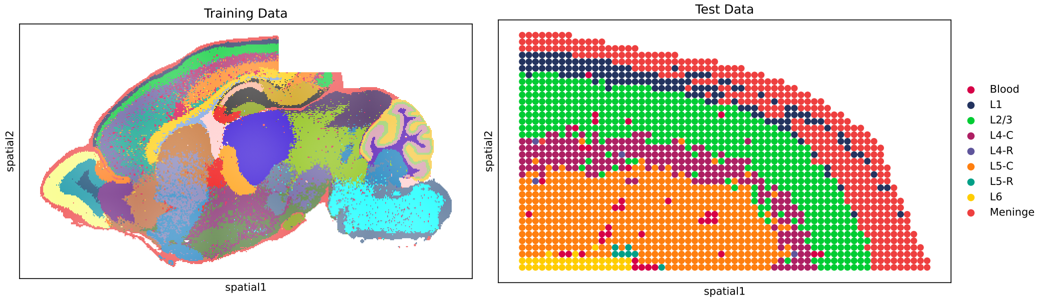


Figure 6.4: Stereo-seq brain experiment setting. We used the fraction of the cortex layer as the test data and used the rest of the spots in the sagittal section as the training data.

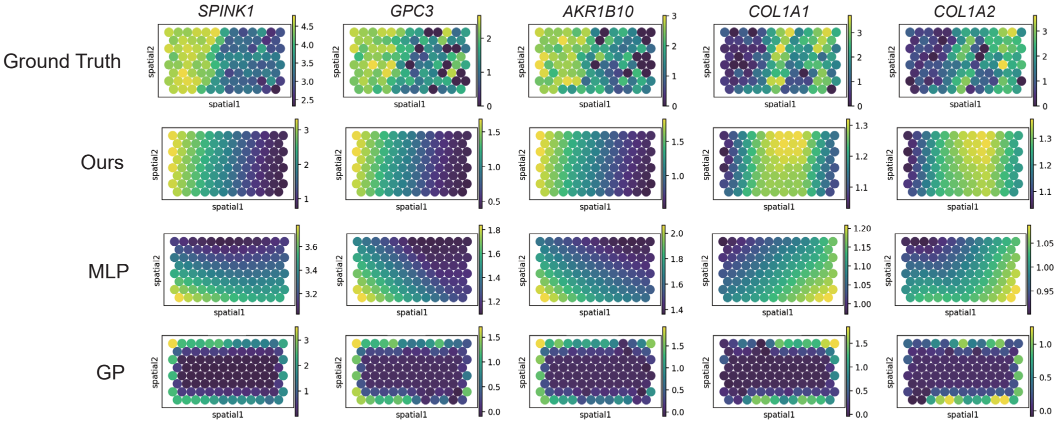


Figure 6.5: Visualization of gene expression predictions of HCC-1L experiment.

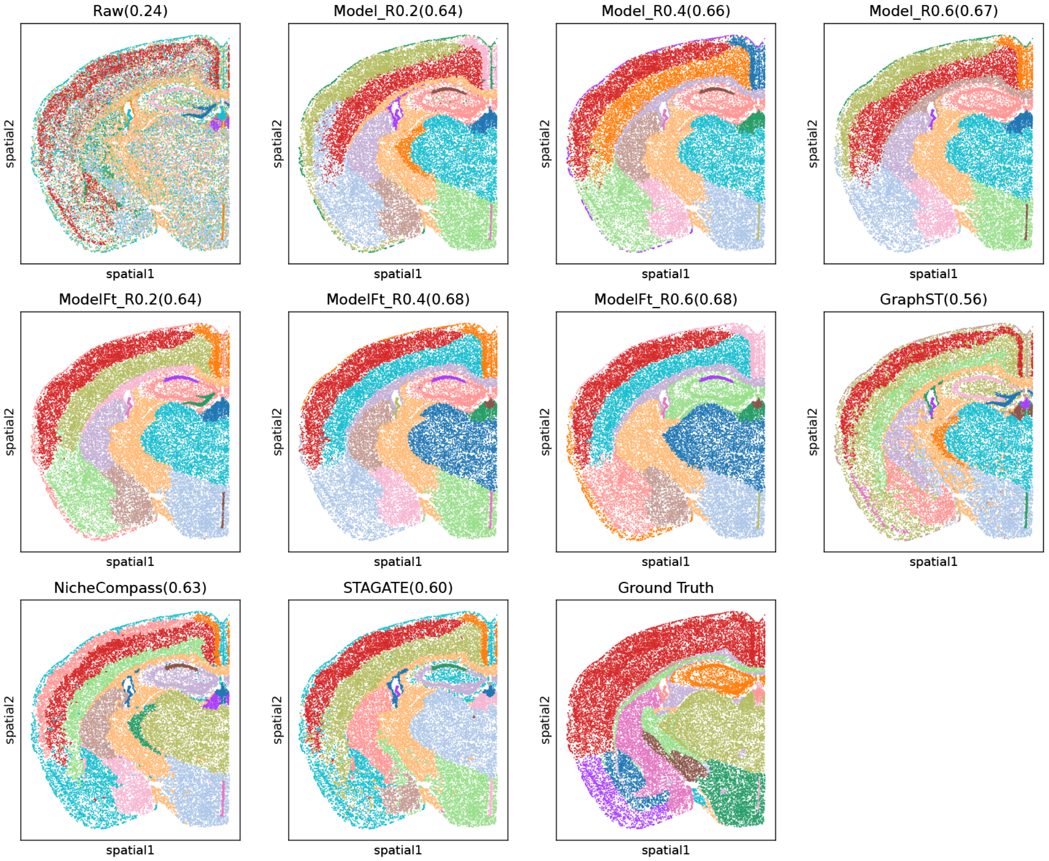


Figure 6.6: Visualization of niche clustering results. Color represents clustered tissue regions at the division level. The number in the subtitle is the adjusted mutual information (AMI) score. 'ModelFt' represents fine-tuned model. '_R0.X' represents the window size of 0.X mm.

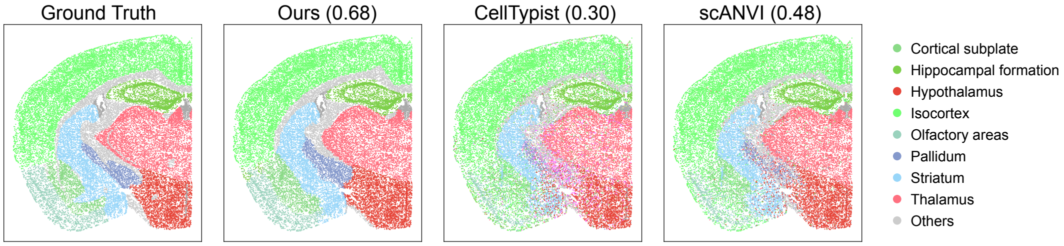


Figure 6.7: Visualization of niche annotation results. Color represents annotated and predicted tissue divisions. The number in the subtitle is the macro F1 score.

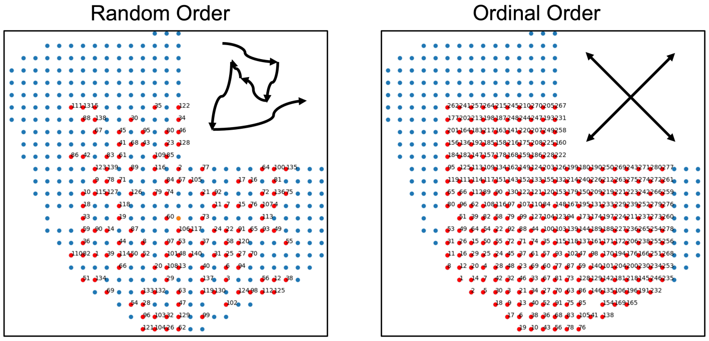


Figure 6.8: Random order and ordinal order in spatial. Given all cells (in blue) in a section, the cells in red constitute a sequence for training. The number next to each cell represents the index in the sequence. In the random serialization strategy, numbers are scattered in space and the cells are not neared. In our proposed ordinal serialization strategy, numbers are sequentially assigned starting from the lower left (smallest x and y values) to the upper right (largest x and y values), but still retaining the randomness.

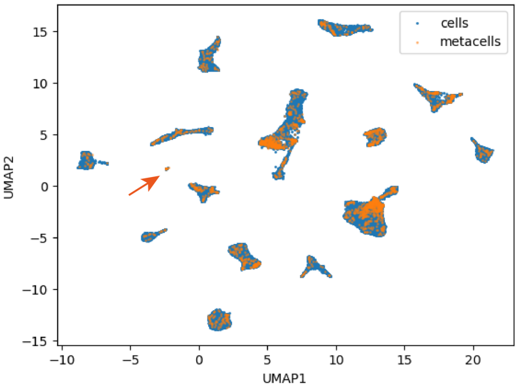


Figure 6.9: UMAP plot of all meta cells and original cells from the PLC dataset. Orange arrow indicates one rare sub-cluster.

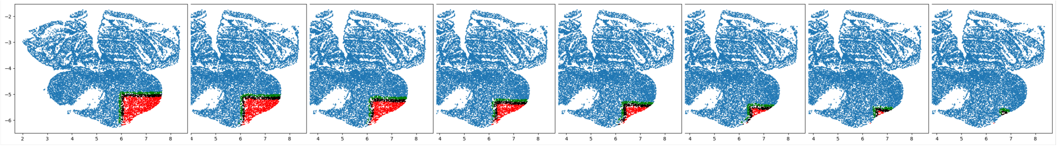

Figure 6.10: Illustration of our iterative generation. Given seen cells (in blue) and spatial locations of unseen cells (in red), our model generated cells iteratively from left to right. For each round, it takes seen cells near the edge of the tissue as the refernce and generates adjacent unseen cells (in black), which will be used as the seen cells in the next generation.

Table 6.1: Full list of differentially expressed genes (DEGs) in PIA_P. (n.s., not significant)

| | Gene name | DEG type (ground truth) | DEG type (prediction) |
|---|---|---|---|
| 1 | *Col5a2* | High | High |
| 2 | *Spp1* | High | High |
| 3 | *Gfap* | High | High |
| 4 | *Ptprc* | High | High |
| 5 | *Lhfpl2* | High | High |
| 6 | *Rnh1* | High | High |
| 7 | *Ctss* | High | High |
| 8 | *Tmem176b* | High | High |
| 9 | *Dcn* | High | High |
| 10 | *Tgfbi* | High | High |
| 11 | *Prkcd* | High | High |
| 12 | *Cldn5* | High | High |
| 13 | *Mdfic* | High | High |
| 14 | *Anxa2* | High | High |
| 15 | *Fn1* | High | High |
| 16 | *Tnc* | High | High |
| 17 | *Ucp2* | High | High |
| 18 | *Maf* | High | High |
| 19 | *Cd44* | High | Low |
| 20 | *Serpinf1* | High | High |
| 21 | *Tmem176a* | High | High |
| 22 | *Cd24a* | High | Low |
| 23 | *Mcm6* | High | Low |
| 24 | *Lsp1* | High | High |
| 25 | *Serpina3n* | High | n.s. |
| 26 | *Col18a1* | High | Low |
| 27 | *Lmo2* | High | High |
| 28 | *Klk6* | High | High |
| 29 | *Cd36* | High | n.s. |
| 30 | *A2m* | High | n.s. |
| 31 | *Penk* | High | n.s. |
| 32 | *Cldn11* | High | High |
| 33 | *Mafb* | High | Low |
| 34 | *Cdkn1a* | High | Low |
| 35 | *Lcp1* | High | Low |
| 36 | *Fyb* | High | n.s. |
| 37 | *Lpl* | High | Low |
| 38 | *Rbp1* | High | High |
| 39 | *Ctsc* | High | Low |
| 40 | *Tgfbr2* | High | High |
| 41 | *Prdm8* | Low | High |
| 42 | *Car4* | Low | High |
| 43 | *Bhlhe22* | Low | High |
| 44 | *Ccn3* | Low | High |
| 45 | *Cnih3* | Low | n.s. |
| 46 | *Krt12* | Low | n.s. |
| 47 | *Slc30a3* | Low | n.s. |
| 48 | *Pvalb* | Low | n.s. |
| 49 | *Chrm1* | Low | High |
| 50 | *Fezf2* | Low | Low |
| 51 | *Kcnj4* | Low | Low |
| 52 | *Tafa1* | Low | Low |
| 53 | *Coro6* | Low | Low |

Continued on next page

Table 6.1 continued from previous page

| | **Gene name** | **DEG type (ground truth)** | **DEG type (prediction)** |
|---|---|---|---|
| 54 | *Rgs6* | Low | Low |
| 55 | *Neurod2* | Low | Low |
| 56 | *Lamp5* | Low | High |
| 57 | *Igfbp6* | Low | Low |
| 58 | *Cpne9* | Low | Low |
| 59 | *Pamr1* | Low | Low |
| 60 | *Bcl11a* | Low | Low |
| 61 | *Adra1b* | Low | Low |
| 62 | *Dkkl1* | Low | n.s. |
| 63 | *Cckbr* | Low | Low |
| 64 | *Chrm3* | Low | Low |
| 65 | *Kcnh3* | Low | High |
| 66 | *Slc17a7* | Low | Low |
| 67 | *Bdnf* | Low | Low |
| 68 | *Myl4* | Low | Low |
| 69 | *Epha4* | Low | Low |
| 70 | *Cbln2* | Low | Low |
| 71 | *Satb2* | Low | Low |
| 72 | *Egr3* | Low | Low |
| 73 | *Hs3st2* | Low | Low |
| 74 | *Pde1a* | Low | Low |
| 75 | *Nwd2* | Low | Low |
| 76 | *Mef2c* | Low | Low |
| 77 | *Rbp4* | Low | Low |
| 78 | *Gabbr2* | Low | Low |
| 79 | *Ldb2* | Low | Low |
| 80 | *Neurod6* | Low | Low |
| 81 | *Fgf13* | Low | Low |
| 82 | *Kcnab3* | Low | Low |
| 83 | *Sv2b* | Low | Low |
| 84 | *Satb1* | Low | Low |
| 85 | *Adcy2* | Low | Low |
| 86 | *Epha10* | Low | Low |
| 87 | *Zmat4* | Low | Low |
